# Unaltered prion disease in mice lacking developmental endothelial locus-1 (Del-1)

**DOI:** 10.1101/348177

**Authors:** Caihong Zhu, Zhihao Li, Bei Li, Manuela Pfammatter, Adriano Aguzzi

## Abstract

Progression of prion diseases is driven by the accumulation of prions in the brain. Ablation of microglia or deletion of the eat-me-signal milk-fat globule EGF factor VIII (Mfge8) accelerate prion pathogenesis, suggesting that microglia defends the brain by phagocytosing prions. Like Mfge8, Developmental endothelial locus-1 (Del-1) is a secreted protein that acts as an opsonin bridging phagocytes and apoptotic cells to facilitate phagocytosis. We therefore asked whether Del-1 might play a role in controlling prion pathogenesis. We first determined the expression pattern of Del-1 in mice, and found that the brain expresses the highest level of Del-1. In mouse brains, Del-1 was mainly expressed by neurons. We then assessed the anti-inflammatory and phagocytosis-promoting functions of Del-1 in prion disease, and determined whether Del-1 complements Mfge8 in prion clearance in mice with a C57BL/6J genetic background. We found that Del-1 deficiency did not change prion disease progression and lesion patterns. Also, prion clearance and PrP^Sc^ deposition were unaltered in Del-1 deficient mice. Additionally, prion-induced neuroinflammation was not affected by Del-1 deficiency. We conclude that Del-1 is not a major determinant of prion pathogenesis in this context.

## 1. Introduction

Prion diseases are lethal neurodegenerative disorders that include scrapie in sheep and goats, bovine spongiform encephalopathy (BSE) in cattle, chronic wasting disease (CWD) in cervids and Creutzfeldt-Jakob disease (CJD) in humans (Aguzzi, et al., 2013b). Prion diseases are characterized by deposition of scrapie prion protein (PrP^Sc^) in the central nervous system (CNS) (Aguzzi and Zhu, 2012). PrP^Sc^ is a misfolded form of the cellular prion protein (PrP^C^), which can seed and convert PrP^C^ into an altered, aggregated conformation. Therefore, prion diseases are transmissible. PrP^Sc^, biochemically defined as a proteinase-K resistant variant of the prion protein, is considered a reliable surrogate marker of the infectious agent. The accumulation of PrP^Sc^ within the central nervous system goes along with astrogliosis, microglial activation, and neuronal loss.

Microglia are the main innate immune cells in the CNS and can defend against various neuroinvasive agents. Removal of microglia by expression of a suicide gene greatly enhances PrP^Sc^ accumulation in prion-infected cultured organotypic cerebellar slices (COCS) and in mice, suggesting that microglia exerts a defensive role against prions (Aguzzi and Zhu, 2017,Falsig, et al., 2008,Zhu, et al., 2016). The prion removal capacity of the brain is considerable, since in *Prnp*-ablated mice, which cannot replicate prions because they lack the conversion substrate, prions are removed within just a few days (Sailer, et al., 1994). However, the defense provided by microglia is only temporary and PrP^Sc^ accumulate over time, eventually leading to prion pathology in wild-type mice.

The molecular mechanisms underlying prion clearance by microglia are not entirely clear. We have reported enhanced prion pathogenesis in mice lacking the astrocytes-borne opsonin protein, milk-fat globule EGF factor VIII (Mfge8). This observation suggests that Mfge8 can opsonize PrP^Sc^ aggregates and facilitate their engulfment by microglia. However, the deficiency of Mfge8 is deleterious only in C57BL/6J × 129Sv, but not in C57BL/6J mice (Kranich, et al., 2010,Kranich, et al., 2008), suggesting the existence of additional factors involved in prion clearance (Aguzzi, et al., 2013a). Triggering receptor expressed on myeloid cells-2 (TREM2), a molecule expressed mainly by microglia in the brain, is important for phagocytosis of apoptotic neurons and suppression of inflammation (Takahashi, et al., 2005). Variants of the TREM2 gene are risk factors for Alzheimer’s disease (Guerreiro, et al., 2013,Jonsson, et al., 2013). This is probably due to impaired phagocytosis and clearance of Aβ (Wang, et al., 2015). However, although TREM2 is involved in prion-induced microglial activation, it is not a main transducer of prion clearance (Zhu, et al., 2015a).

Developmental endothelial locus-1 (Del-1) was originally found as an endothelial cell-derived extracellular matrix protein structurally containing three EGF-like repeats and two discoidin I-like domains. Del-1 regulates vascular morphogenesis or remodeling (Hidai, et al., 1998). Del-1 has also emerged as an inhibitor of leukocyte-endothelial interaction by interfering with lymphocyte function-associated antigen (LFA)-1-dependent leukocyte recruitment (Choi, et al., 2008), thereby limiting ischemia-related angiogenesis (Klotzsche-von Ameln, et al., 2017). Recently, Del-1 has appeared as a potent antagonist of interleukin 17 (IL-17)-triggered neutrophil-mediated inflammatory bone loss and osteoclast-induced bone resorption (Eskan, et al., 2012,Maekawa, et al., 2015,Shin, et al., 2015). The anti-inflammatory function of Del-1 has been observed also in the CNS, and Del-1^−/−^ mice develop accelerated experimental autoimmune encephalomyelitis (EAE) pathogenesis due to increased neuroinflammation and demyelination (Choi, et al., 2015). Another study found that Del-1 is expressed by various cells in bone marrow and functions as a regulator of myelopoiesis in the hematopoietic stem cell (HSC) niche (Mitroulis, et al., 2017). As a structural homologue of Mfge8, Del-1 can also bind to α_v_β_3_ integrin on phagocytes via an Arg-Gly-Asp (RGD) motif in the second EGF domain, and to phosphatidylserine exposed on the outer leaflet of apoptotic cells through its discoidin I-like domains, therefore bridging apoptotic cells and phagocytes to facilitate uptake and removal of apoptotic cells (Hanayama, et al., 2004). However, it has also been reported that Del-1 attenuates complement-dependent phagocytosis by blocking macrophage-1 antigen (Mac-1) integrin (Mitroulis, et al., 2014).

In this study, we aimed to investigate whether the phagocytosis-promoting and anti-inflammatory functions of Del-1 may have an impact onto prion pathogenesis, and to determine whether Del-1 complements Mfge8 in prion clearance in mice with a C57BL/6J genetic background. We first determined the expression pattern of Del-1 and found that Del-1 is mainly expressed by neurons in mouse brains. We observed that Del-1^−/−^ mice displayed disease progression and lesion patterns similar to those of their heterozygous (Del-1^+/−^) and wild-type (Del-1^+/+^) littermates. Importantly, PrP^Sc^ deposition was not altered by Del-1 deficiency, suggesting that Del-1 is not involved in prion clearance in vivo. Furthermore, Del-1 deficiency did not overtly affect prion-induced neuroinflammation. On the basis of these observations, we conclude with a high degree of confidence that Del-1 is not a major determinant of prion clearance and does not impact prion pathogenesis in any measurable way, Del-1 does not complement Mfge8 in prion clearance in mice with a C57BL/6J genetic background.

## 2. Material and methods

### 2.1 Ethical statement

All animal experiments were carried out in strict accordance with the Rules and Regulations for the Protection of Animal Rights (Tierschutzgesetz and Tierschutzverordnung) of the Swiss Bundesamt für Lebensmittelsicherheit und Veterinärwesen and were preemptively approved by the Animal Welfare Committee of the Canton of Zürich (permit # 41/2012).

### 2.2 Animals

Del-1^−/−^ mice with a C57BL/6J genetic background (Choi, et al., 2008) were generated by a targeted replacement of exon 1 by a reporter gene β-galactosidase, therefore expressing β-galactosidase instead of Del-1 under the control of the endogenous Del-1 promoter. Del-1^−/−^ mice were backcrossed again to C57BL/6J mice to obtain Del-1^+/−^ offspring, which were then intercrossed to generate Del-1^+/+^ (wild type), Del-1^+/−^ and Del-1^−/−^ mice for experiments described here. All animals were maintained in a high hygienic grade facility under a 12 h light/12 h dark cycle (from 7 am to 7 pm) at 21±1°C, and fed with diet and water ad libitum.

### 2.3 Prion inoculation

Mice at 6-8 weeks old were intracerebrally (i.c) inoculated with 30 μl of brain homogenate diluted in PBS with 5% BSA and containing 3 × 10^5^ LD50 units of the Rocky Mountain Laboratories scrapie strain (passage 6, thus called RML6). Mice were monitored and actions were taken to minimize animal suffering and distress according to details described previously (Zhu, et al., 2015b). Scrapie was diagnosed according to clinical criteria (ataxia, limb weakness, front leg paresis and rolling). Mice were sacrificed by CO_2_ inhalation on the day of appearance of terminal clinical signs of scrapie (specific criteria referred to (Zhu, et al., 2015b)), organs were taken and then were either snap-frozen for biochemical analysis or fixed in 4% formalin for histological assessment. The time elapsed from prion inoculation to the terminal stage of disease was defined as incubation time for the survival study.

### 2.4 Quantitative real-time PCR (qRT-PCR)

Total RNA was extracted using TRIzol (Invitrogen Life Technologies) according to the manufacturer’s instruction. The quality of RNA was analyzed by Bioanalyzer 2100 (Agilent Technologies), RNAs with RIN>7 were used for cDNA synthesis. cDNAs were synthesized from ~1 μg total RNA using QuantiTect Reverse Transcription kit (QIAGEN) according to the manufacturer’s instruction. Quantitative real-time PCR (qRT-PCR) was performed using the SYBR Green PCR Master Mix (Roche) on a ViiA7 Real-Time PCR system (Applied Biosystems). The following primer pairs were used: GAPDH sense, 5´-CCA CCC CAG CAA GGA GAC T-3´; antisense, 5´-GAA ATT GTG AGG GAG ATG CT-3´. Del-1 sense, 5´-GCC TGG CTT TTG GTT GGA CT-3´; antisense, 5´-ACC TGC GAA GCC TTC TGG AC-3´. Mfge8 sense, 5´-ATA TGG GTT TCA TGG GCT TG-3´; antisense, 5´-GAG GCT GTA AGC CAC CTT GA-3´. TNFα sense, 5´-CAT CTT CTC AAA ATT CGA GTG ACA A-3´; antisense, 5´-TGG GAG TAG ACA AGG TAC AAC CC-3´. IL-1β sense, 5´-CAA CCA ACA AGT GAT ATT CTC CAT G-3´; antisense, 5´-GAT CCA CAC TCT CCA GCT GCA-3´. IL-6 sense, 5´-TCC AAT GCT CTC CTA ACA GAT AAG-3´; antisense, 5´-CAA GAT GAA TTG GAT GGT CTT G-3´.

Expression levels were normalized using GAPDH.

### 2.5 Immunohistochemistry

For X-gal (5-bromo-4-chloro-3-indolyl-β-d-galactopyranoside) staining, cryosections were fixed at 0.4% glutaraldehyde at 4°C for 5 min. After washing with PBS, sections were pretreated with permeabilization buffer (PBS containing 0.02% NP-40, 0.01% sodium deoxycholate and 2 nM MgCl2) at RT for 30 min. Sections were washed with PBS and incubated with staining solution (5 mM FeKCN(II), 5 mM FeKCN(III), 2 mM MgCl2 and 1mg/ml X-Gal (Promega V3941)) at 37°C for 24 hours. Sections were washed again with PBS containing 0.02% Igepal and 2 mM MgCl2, and mounted with DAKO faramount. Only cells that express β-galactosidase can catalyze the hydrolysis of X-gal into galactose and 5-bromo-4-chloro-3-hydroxyindole, the second compound is then oxidized into 5,5’-dibromo-4,4’-dichloro-indigo that will visualize the cells blue. Sections were imaged using a Zeiss Axiophot light microscope.

Prion-infected brain tissues were harvested and fixed in formalin, followed by treatment with concentrated formic acid for 60 min to inactivate prion infectivity and embedded in paraffin. Paraffin sections (2 μm) of brains were stained with hematoxylin/eosin (HE) to visualize prion-induced lesions and vacuolation. For the histological detection of partially proteinase K-resistant prion protein deposition, deparaffinized sections were pretreated with formaldehyde for 30 min and 98% formic acid for 6 min, and then washed in distilled water for 30 min. Sections were incubated in Ventana buffer and stains were performed on a NEXEX immunohistochemistry robot (Ventana instruments, Switzerland) using an IVIEW DAB Detection Kit (Ventana). After incubation with protease 1 (Ventana) for 16 min, sections were incubated with anti-PrP SAF-84 (1:200, SPI bio, A03208) for 32 min. Sections were counterstained with hematoxylin. To detect astrogliosis and microglial activation, brain sections were deparaffinized through graded alcohols, anti-GFAP antibody (1:300; DAKO, Carpinteria, CA) were applied for astrogliosis, anti-Iba1 antibody (1:1000; Wako Chemicals GmbH, Germany) was used for highlighting activated microglial cells. Stainings were visualized using DAB (Sigma-Aldrich), hematoxylin counterstain was subsequently applied. Sections were imaged using a Zeiss Axiophot light microscope.

### 2.6 Immunofluorescent staining

Cryosections were first fixed in acetone at room temperature for 10 min, followed by blocking in blocking buffer (0.5% BSA (Albumin Fraktion V, Roth, Karlsruhe), 1% goat serum (X0907, DAKO) and 1% Triton X-100 (T9284-100ML, Sigma-Aldrich) in PBS) for 30 min. Sections were then incubated with primary antibodies diluted in staining buffer (0.5% BSA, PBS, 1% goat serum and 0.1% Triton X) at 4°C overnight. After washing, sections were incubated with secondary antibodies conjugated with different fluorophores at room temperature for 1 hour. After washing, nuclei were stained with DAPI (sigma, D9542). Sections were mounted with DAKO Faramount Aqueous mounting medium. Images were captured using confocal laser scanning microscope FluoView Fv10i (Olympus). The following antibodies were used for immunofluorescent staining: Anti-β-galactosidase (1:500, Abcam, ab9361); Anti-MAP2 (1:300, Abcam, ab32454); Anti-GFAP (1:500, DAKO, Z0334); Anti-CD68 (1:200, Bio-Rad, MCA1957); anti-MAG (1:200, Millipore, MAB1567); DyLight 488 goat anti chicken (1:500, Abcam, ab96947); Alexa Fluor 594 Goat anti rabbit (1:500, Invitrogen, A11037); Alexa Fluor 594 goat anti rat (1:500, Invitrogen, A11007); Alexa Fluor 594 goat anti mouse (1:500, Invitrogen, A11032).

### 2.7 Western blot analysis

To detect PrP^C^ in the mouse brains, one hemisphere of each brain was homogenized with buffer PBS containing 0.5% Nonidet P-40 and 0.5%CHAPSO. Total protein concentration was determined using the bicinchoninic acid assay (Pierce). ~8 ug protein was loaded and separated on a 12% Bis-Tris polyacrylamide gel (NuPAGE, Invitrogen) and then blotted onto a nitrocellulose membrane. Membranes were blocked with 5% (wt/vol) Topblock (LuBioScience) in PBS supplemented with 0.1% Tween 20 (vol/vol) and incubated with primary antibodies POM1 in 1% Topblock (400 ng ml^−1^) overnight. After washing, the membranes were then incubated with secondary antibody horseradish peroxidase (HRP)-conjugated goat anti-mouse IgG (1:10,000, Jackson ImmunoResearch, 115-035-003). Blots were developed using Luminata Crescendo Western HRP substrate (Millipore) and visualized using the Stella system (model 3000, Bio-Rad). To avoid variation in loading, the same blots were stripped and incubated with anti-actin antibody (1:10,000, Millipore). The PrP^C^ signals were normalized to actin as a loading control. To detect Mfge8 protein in mouse brains by Western blot, 20 μg of total brain protein were loaded and anti-Mfge8 antibody (1:1000; R&D Systems, AF2805) and horseradish peroxidase (HRP)-conjugated donkey anti-goat IgG(1:10,000, Jackson ImmunoResearch, 705-035-147) were used as primary and secondary antibodies, respectively. Actin was used as the loading control.

To detect PrP^Sc^, prion infected forebrains were homogenized in sterile 0.32 M sucrose in PBS. Total protein concentration was determined using the bicinchoninic acid assay (Pierce). Samples were adjusted to 20 μg protein in 20 μl and digested with 25 μg ml^−1^ proteinase K for 30 min at 37°C. PK digestion was stopped by adding loading buffer (Invitrogen) and boiling samples at 95°C for 5 min. Proteins were then separated on a 12% Bis-Tris polyacrylamide gel (NuPAGE, Invitrogen) and blotted onto a nitrocellulose membrane. POM1 and horseradish peroxidase (HRP)-conjugated goat anti-mouse IgG were used as primary and secondary antibodies, respectively. Blots were developed using Luminata Crescendo Western HRP substrate (Millipore) and visualized using the FUJIFILM LAS-3000 system. To detect Iba1 and GFAP in prion-infected brains by Western blot, 20 μg of total brain protein were loaded and anti-Iba1 antibody (1:1000; Wako Chemicals GmbH, Germany, 019-19741), anti-GFAP antibody (D1F4Q) XP Rabbit mAb (1:3000; Cell Signaling Technology, 12389) and horseradish peroxidase (HRP)-conjugated goat anti-rabbit IgG (1:10,000, Jackson ImmunoResearch, 111-035-045) were used as primary and secondary antibodies, respectively. Actin was used as the loading control.

### 2.8 Real-time quaking-induced conversion assay (RT-QuIC)

RT-QuIC assays of prion-infected mouse brain homogenates were performed as previously described (Frontzek, et al., 2016). Briefly, recombinant hamster full-length (23–231) PrP (HaPrP) was expressed in Rosetta2(DE3)pLysS *E.coli* competent cells and purified by affinity chromatography using Ni^2+^-nitrilotriacetic acid Superflow resin (QIAGEN). In the RT-QuIC assay, recombinant HaPrP was used as substrate for PrP^Sc^-catalyzed conversion. RT-QuIC reactions containing HaPrP substrate protein at a final concentration of 0.1 mg mL^−1^ in PBS (pH 7.4), 170 mM NaCl, 10 μM EDTA, 10 μM Thioflavin T were seeded with 2 μL of serially diluted brain homogenates in a total reaction volume of 100 μL. NBH- and RML6-treated brain homogenates were used as negative and positive controls, respectively. The RT-QuIC reactions were amplified at 42 °C for 100 h with intermittent shaking cycles of 90 s shaking at 900 rpm in double orbital mode and 30 s resting using a FLUOstar Omega microplate reader (BMG Labtech). Aggregate formation was followed by measuring the thioflavin T fluorescence every 15 min (450 nm excitation, 480 nm emission; bottom read mode).

### 2.9 Statistical analysis

Results are presented as the mean of replicas ± standard error of the mean (SEM). An unpaired Student’s t-Test was used to assess statistical significance between two experimental groups. One-way ANOVA with Tukey’s multiple comparison test or Dunnett’s multiple comparison test was used for comparison of all columns to a control column for statistical analysis of experiments involving the comparison of three or more groups. For prion inoculation experiments, incubation times were analyzed using the Kaplan-Meier method and compared between groups using Log-rank (Mantel-Cox) test. p-values <0.05 were considered statistically significant.

## 3. Results

### 3.1 Del-1 is highly expressed in the mouse brain in an age-dependent manner

Del-1 is a secreted glycoprotein. We have not been able to acquire a specific antibody against the mouse Del-1 protein, and two commercially available antibodies (Novus Biologicals, NBP1-28632; Proteintech, 12580-1-AP) were found to only yield nonspecific bands in Western blots (Supplemental figure 1). Therefore, we assessed Del-1 expression using quantitative reverse-transcription PCR (qRT-PCR). We first compared expression levels of Del-1 in brain, liver, thymus, spleen, lung, kidney, heart, limb muscle and bone marrow collected from 2-month old wild-type C57BL/6J mice. Organs from Del-1^−/−^ littermates were also collected and used as negative controls. Consistent with a previous report (Choi, et al., 2008), we found that mouse brains expressed the highest level of Del-1 of all organs; the lungs also expressed conspicuous levels (Figure 1A). However, the expression of Del-1 was barely detectable in other organs (Figure 1A). In contrast, Mfge8, the structural and functional homologue of Del-1 produced by follicular dendritic cells and astrocytes, showed ubiquitous expression among different organs with the highest expression in secondary lymphoid organs such as spleen (Figure 1B). The restricted expression pattern of Del-1 suggested that Del-1 may exert certain specific functions in the central nervous system.

**Figure 1:**
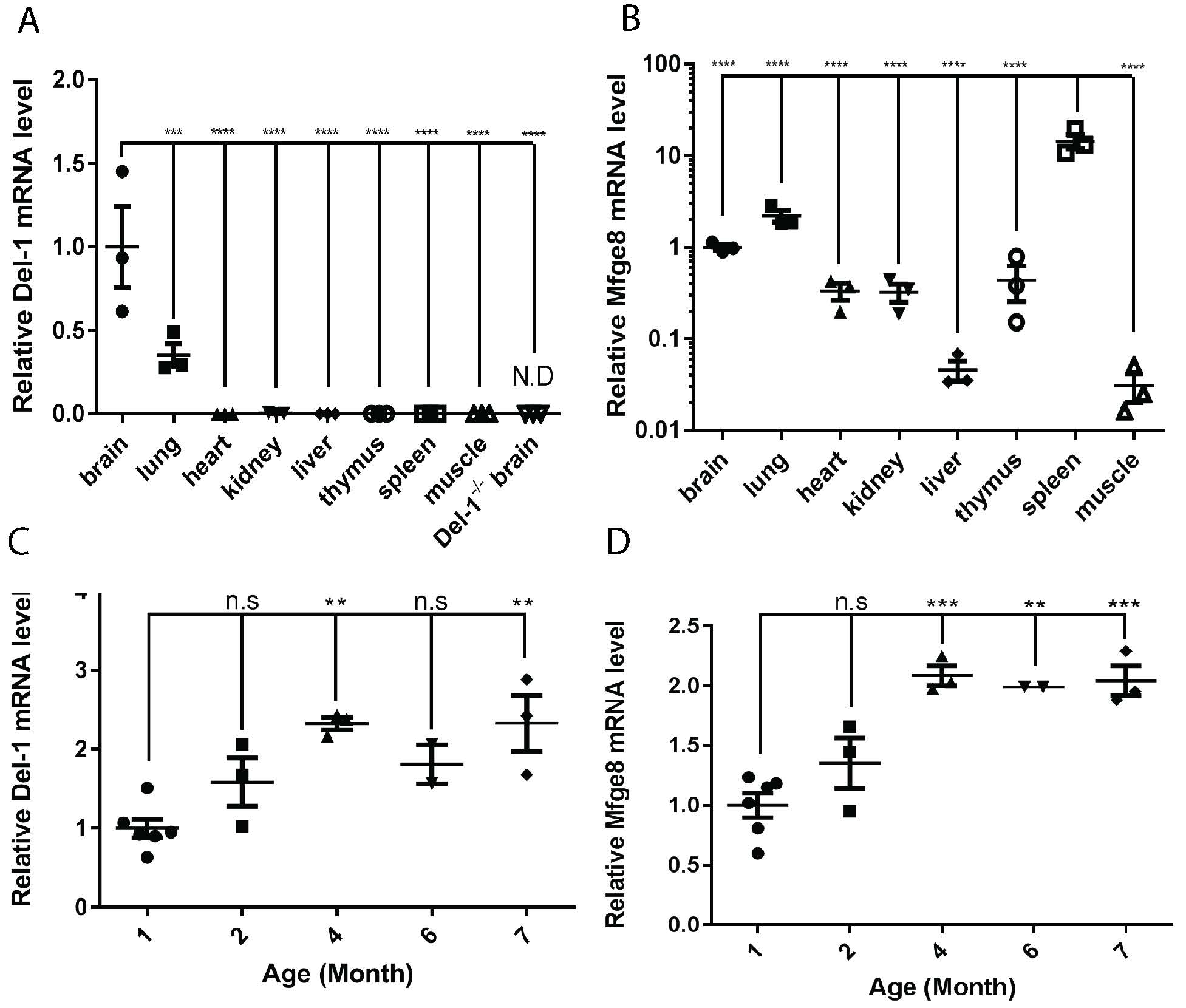
(A-B) qRT-PCR of Del-1 (A) and Mfge8 (B) expression in various mouse organs. Del-1 was significantly expressed in murine brain and lungs, but was barely detectable in other organs. In contrast, Mfge8 was ubiquitously expressed in various organs, with the spleen expresses the highest level. ND: not detectable. ***, p<0.001; ****, p<0.0001. (C-D) qRT-PCR of Del-1 (C) and Mfge8 (D) expression in mouse brains at different ages. Both Del-1 and Mfge8 expression increased from 1-month to 4-months old and stabilized thereafter. **, p<0.01; ***, p<0.001; n.s, not significant.

The observed expression of Del-1 during the early embryogenesis stage corresponded a developmentally restricted pattern (Hidai, et al., 1998). To determine whether Del-1 expression in adult brain is also developmentally regulated, we carried out qRT-PCR on mRNA isolated from C57BL/6J mice at the age of 1, 2, 4, 6, and 7 months. We observed that Del-1 expression in the brain increased from 1-month to 4-month old and then remained constant thereafter (Figure 1C). We also analyzed the expression of Mfge8 in the brain at different ages. Interestingly, Mfge8 showed a similar expression pattern to Del-1, with the expression level of Mfge8 increasing steadily from 1-month to 4-months old followed by a stabilization at 7-months old (Figure 1D).

### 3.2 Cerebral Del-1 is mainly expressed by neurons

Del-1 has been reported to be expressed by endothelial cells and a subset of macrophages, as well as several cells in hematopoietic stem cell niche (Mitroulis, et al., 2017). However, a detailed study on the expression pattern of Del-1 in the CNS has not been performed. To determine which cell types in the brain express Del-1, we applied immunohistochemical staining to wild-type C57BL/6J mouse brain sections. However, we could not detect Del-1 specific signal immunolabeling in wild-type mice, since a similar signal was also observed in Del-1^−/−^ brain sections (data not shown). We then analyzed Del-1^−/−^ mice in which the exon1 of Del-1 open reading frame was replaced by the β-galactosidase expression cassette. Expression of β-galactosidase is under the control of the endogenous Del-1 promoter, therefore the expression pattern of β-galactosidase could indicate the expression pattern of endogenous Del-1. We then first performed an X-gal (5-bromo-4-chloro-3-indolyl-β-d-galactopyranoside) staining on brain sections prepared from Del-1^−/−^ mice to visualize cells expressing β-galactosidase. Sections from wild type littermates were used as negative controls. We observed intense blue staining in the granular layer of the cerebellum and dentate gyrus of the hippocampus in Del-1^−/−^ brain sections (Figure 2A), suggesting that Del-1 promoter is highly active in granular neurons. These results implicate that granule neurons are the main cells expressing endogenous Del-1.

**Figure 2:**
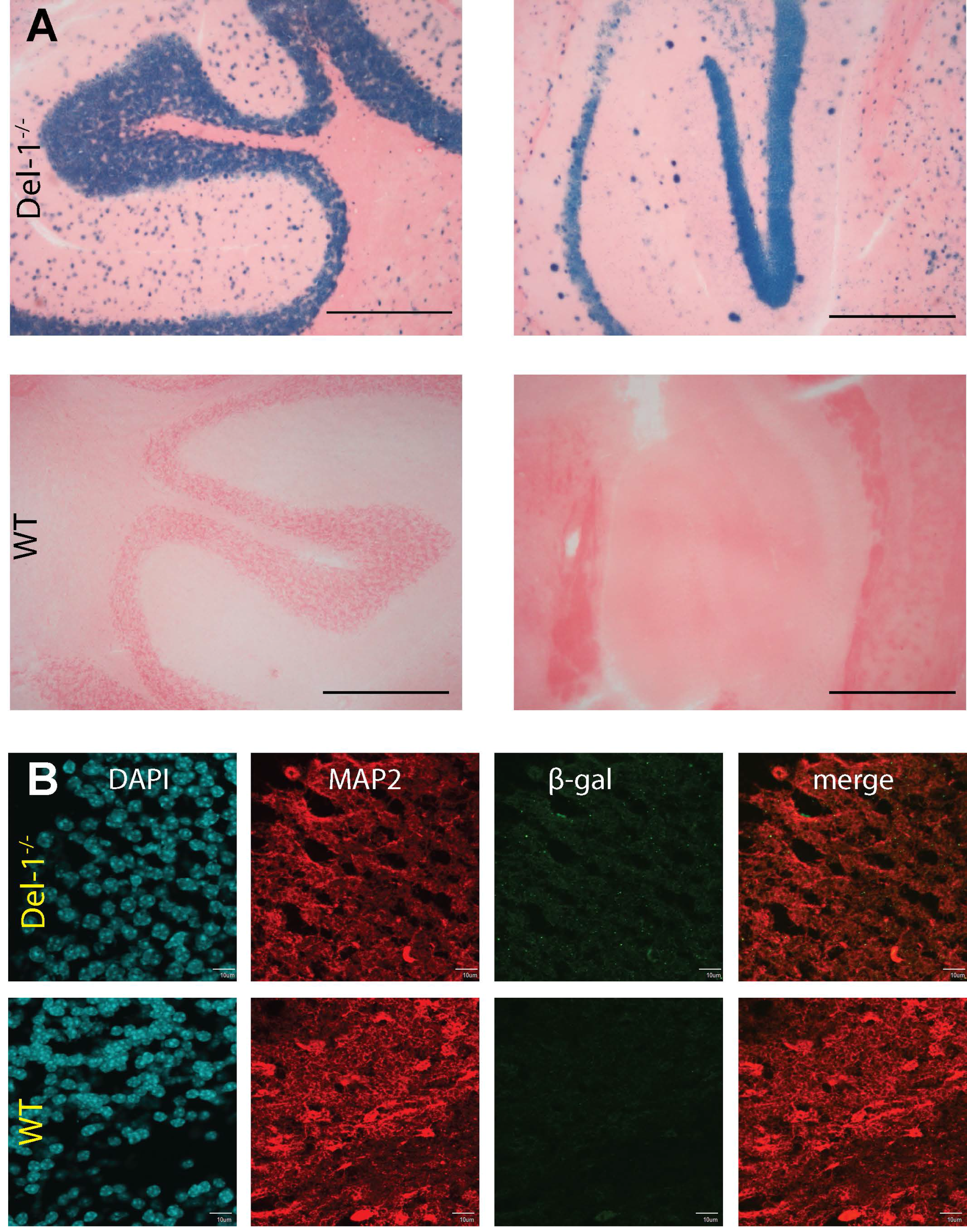
(A) X-gal staining on Del-1^−/−^ and wild type mouse brains. X-gal staining detected intense signals in the granular layer of the cerebellum (left panel) and hippocampus (right panel) in Del-1^−/−^ mouse brains. Whereas in the negative control wild type mouse brains, no X-gal staining was detected. Scale bars: 200um. (B) Immunofluorescent co-staining of β-galactosidase and neuronal markers MAP2 on Del-1^−/−^ and wild type mouse brains. Colocalization of β-galactosidase stain (green dots) and MAP2 was observed in Del-1^−/−^ mouse brains (upper panel). No β-galactosidase stain (green dots) was detected in negative control wild type mouse brains (lower panel) Scale bars: 10um.

To confirm the above observation that neurons are the primary cell type expressing Del-1, we performed immunofluorescent co-stainings using antibodies against β-galactosidase and specific cell markers. Brain tissues from Del-1^−/−^ and C57BL/6J wild type mice were co-stained with anti-β-galactosidase antibodies and one of the cell markers: MAP2 for neurons, GFAP for astrocytes, CD68 for microglia, and MAG for oligodendrocytes. Green dots representing β-galactosidase were observed in Del-1^−/−^ but not in wild-type brain sections (Figure 2B). The β-galactosidase signal predominantly co-localized with the neuronal cell marker MAP2 (Figure 2B), suggesting that β-galactosidase was mainly expressed in neurons of Del-1^−/−^ mice. No colocalization was observed between β-galactosidase and GFAP, CD68 or MAG (Supplemental figure 2), suggesting that astrocytes, microglia and oligodendrocytes do not express Del-1. Our observation not only confirmed previous investigations but also expanded what was reported by another study (Choi, et al., 2015). We therefore conclude that Del-1 is mainly expressed by neurons in mouse brains.

### 3.3 Del-1 deficiency does not affect prion progression and lesion pattern

Del-1 deficiency did not change the transcription of *Prnp* and the protein levels of PrP^C^ in brains (Figure 3A-B), which are major determinants of susceptibility to prion diseases. To determine the role of Del-1 in prion diseases, we tested whether Del-1 deficiency could affect prion disease progression and prion-induced lesion pattern in mouse brains. We then intracerebrally inoculated 30 μl of diluted infectious brain homogenate, corresponding to 3 × 10^5^ LD_50_ units of RML6 prions (a prion strain derived from the Rocky Mountain Laboratory, passage 6) into Del-1^+/+^, Del-1^+/−^, and Del-1^−/−^ littermates. Prion-inoculated mice were monitored every other day for signs of scrapie, and mice were sacrificed when they reached the terminal stage of disease. Incubation times were calculated by the time elapsed from prion inoculation to terminal stage. We observed that Del-1^+/+^, Del-1^+/−^ and Del-1^−/−^ mice succumbed to prion disease in a similar rate (median survival: 186 dpi for female Del-1^+/+^ mice (n=6), 191.5 dpi for female Del-1^+/−^(n=16) and 193.5 dpi for female Del-1^−/−^ mice (n=8), p=0.26; median survival 196.5 dpi for male Del-1^+/+^ mice(n=14), 195 dpi for male Del-1^+/−^(n=19) and 197 dpi for male Del-1^−/−^ mice (n=11), p=0.87) (Figure 3C). These results indicate that Del-1 deficiency does not significantly affect prion progression.

We then analyzed the histology of brain sections prepared from prion-infected terminally sick Del-1^+/+^, Del-1^+/−^ and Del-1^−/−^ mice. The typical histological features of prion disease, including spongiform vacuolation, were observed in all mice with different genotypes (Figure 3D). Lesion pattern analysis failed to find any qualitative differences between the three groups (Figure 3D). These results indicate that Del-1 deficiency does not affect prion-induced lesion pattern in mouse brains.

**Figure 3:**
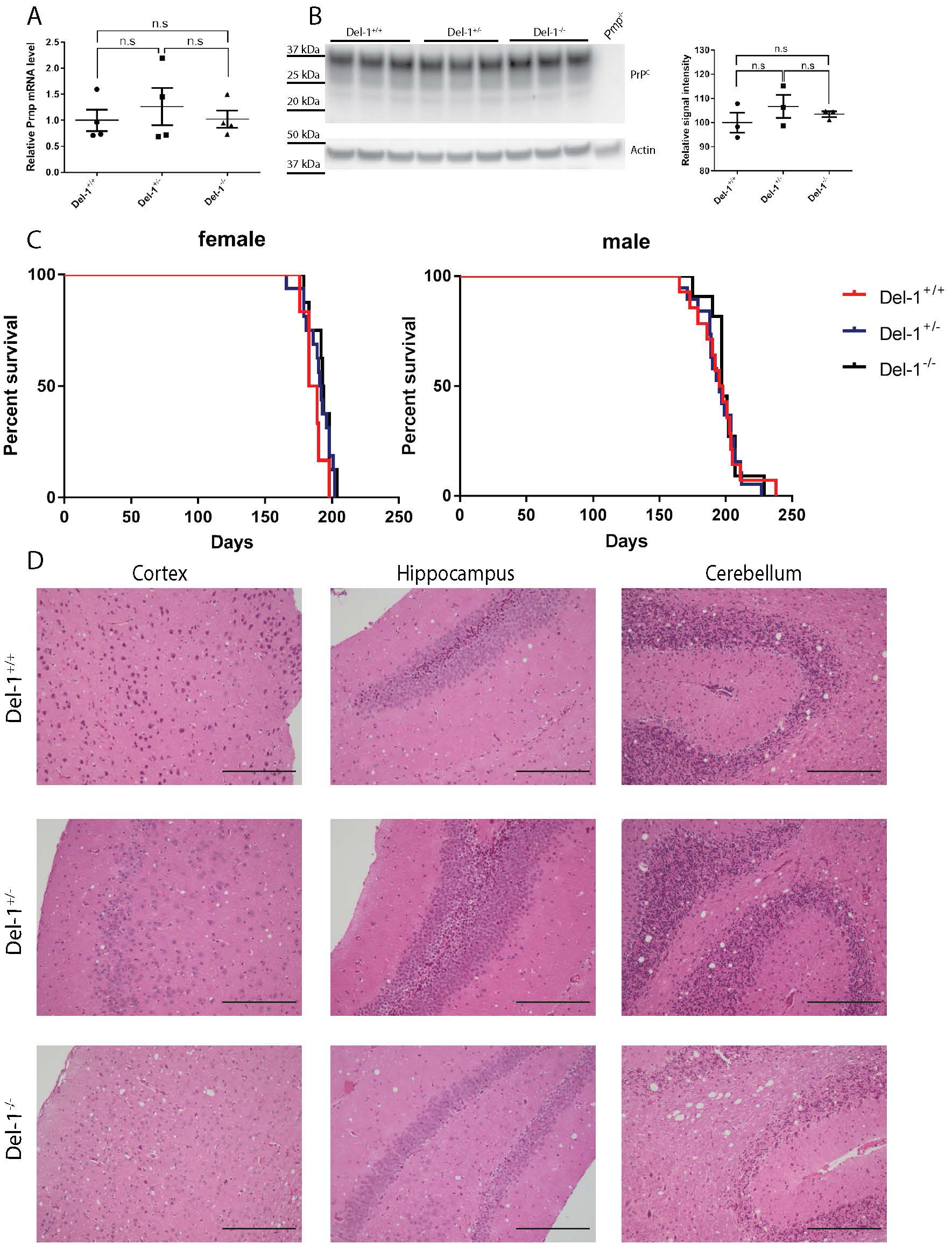
(A-B) *Prnp* qRT-PCR (A) and PrP^C^ Western blot (B left; densitometric quantification of the Western blot in B right) in Del-1^+/+^ (WT), Del-1^+/−^ and Del-1^−/−^ mouse brains. No significant difference of *Prnp* mRNA and PrP^C^ protein was observed between Del-1^+/+^ (WT), Del-1^+/−^ and Del-1^−/−^ mouse brains. (C) Survival curve of Del-1^+/+^ (WT), Del-1^+/−^ and Del-1^−/−^ littermates intracerebrally inoculated with RML6. There was no significant difference between three groups in both genders (n=6~19, n.s *p*>0.05). (D) Representative histology of terminally sick mouse brains from Del-1^+/+^ (WT), Del-1^+/−^ and Del-1^−/−^ littermates stained for hematoxylin and eosin (H&E). There was no obvious difference between three groups in vacuolation and lesion pattern in cortex, hippocampus and cerebellum, Scale bars: 200um.

### 3.4 Del-1 deficiency does not alter PrP^Sc^ accumulation

We next sought to determine whether Del-1, which is similar to its structural and functional homologue Mfge8, is involved in prion clearance in mouse brains. If Del-1 were to contribute to prion clearance, Del-1^−/−^ mice would display more PrP^Sc^ deposition in their brains. We performed PrP^Sc^ staining on brain sections prepared from prion-infected terminally sick Del-1^+/+^, Del-1^+/−^ and Del-1^−/−^ mice. However, we observed similar PrP^Sc^ deposition level in all three groups (Figure 4A). We next carried out Western blot to detect proteinase K-resistant PrP^Sc^ in brains of prion-infected terminally sick Del-1^+/+^, Del-1^+/−^ and Del-1^−/−^ mice. Again, we found similar level of PrP^Sc^ in mouse brains of the three genotypes (Figure 4B). Furthermore, we assessed the seeding activity in brain homogenates of prion-infected terminally sick Del-1^+/+^, Del-1^+/−^ and Del-1^−/−^ mice by RT-QuIC. We detected a similar prion 50% seeding dose (SD50) in all three groups (Figure 4C). These results suggest that, in contrast to Mfge8, Del-1 does not contribute to prion clearance and PrP^Sc^ accumulation in mouse brains.

**Figure 4:**
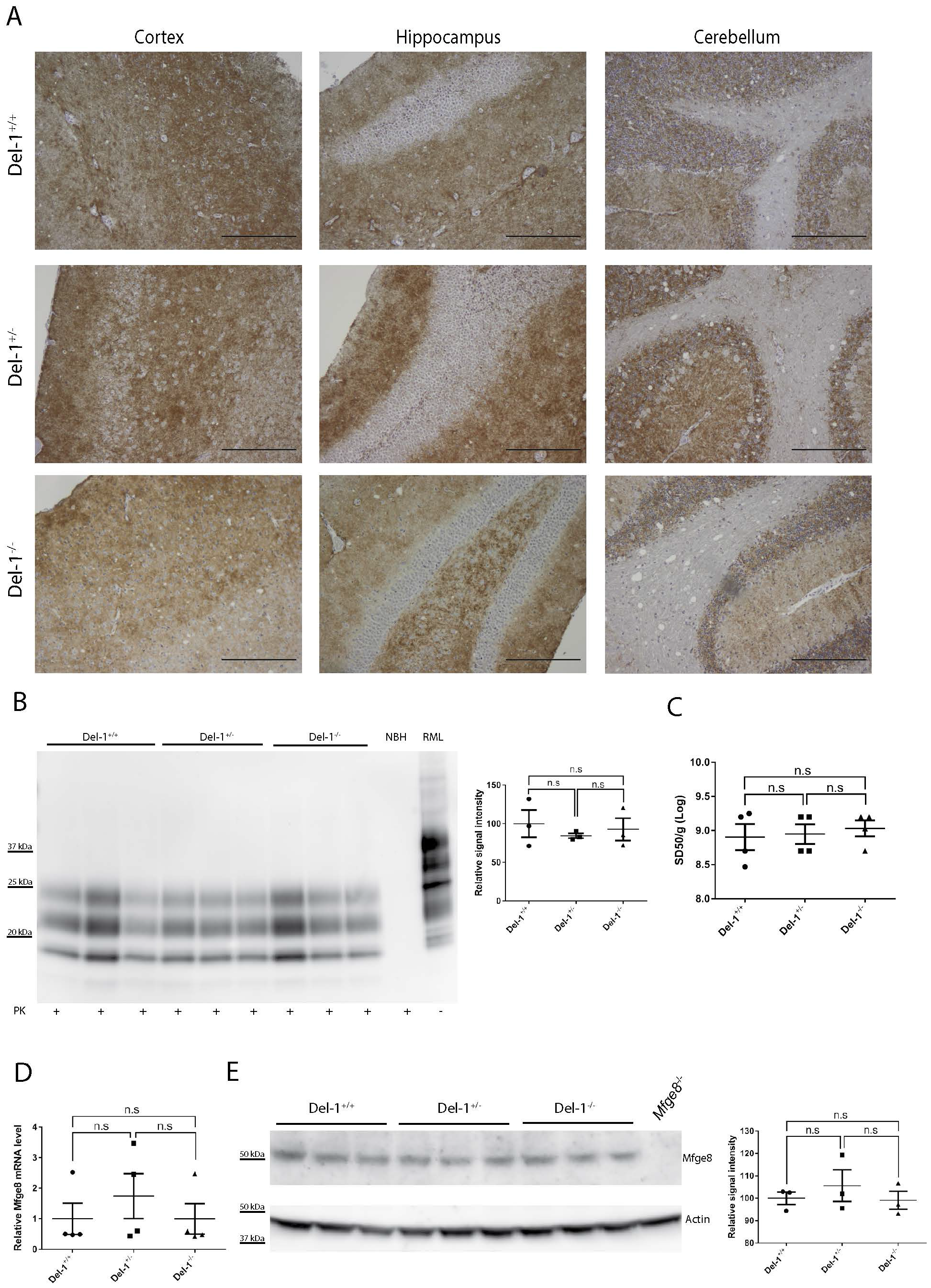
(A) Representative histology of terminally sick mouse brains from Del-1^+/+^ (WT), Del-1^+/−^ and Del-1^−/−^ littermates stained for SAF84. There was no obvious difference between three groups in PrP^Sc^ deposition in cortex, hippocampus and cerebellum, Scale bars: 200um. (B) Left: Western blot for proteinase K resistant PrP^Sc^ in terminally sick mouse brains. Right: densitometric quantification of the Western blot showed no significantly difference between Del-1^+/+^ (WT), Del-1^+/−^ and Del-1^−/−^ littermates (n=3, n.s *p*>0.05). NBH: non-infectious brain homogenates. (C) RT-QuIC assay of terminally sick mouse brains from Del-1^+/+^ (WT), Del-1^+/−^ and Del-1^−/−^ littermates showed similar level of 50% prion seeding dose in these three groups (n=4, n.s *p*>0.05). (D-E) *Mfge8* qRT-PCR (D) and Mfge8 Western blot (E left; densitometric quantification of the Western blot in E right) in Del-1^+/+^ (WT), Del-1^+/−^ and Del-1^−/−^ mouse brains. No significant difference of *Mfge8* mRNA and Mfge8 protein was observed between Del-1^+/+^ (WT), Del-1^+/−^ and Del-1^−/−^ mouse brains.

To test the possibility that Del-1 deficiency could upregulate Mfge8 expression through a compensatory mechanism, therefore blunting the acceleration of prion progression in Del-1^−/−^ mice, we assessed the expression of Mfge8 in Del-1^+/+^, Del-1^+/−^ and Del-1^−/−^ mouse brains. Both qRT-PCR and Western blot failed to detect obvious change of Mfge8 mRNA and protein levels in Del-1^−/−^ mouse brains (Figure 4D-E), suggesting that the unchanged prion pathogenesis in Del-1^−/−^ mice is not due to a compensatory effect of its homologue Mfge8.

### 3.5 Del-1 deficiency does not affect prion-induced neuroinflammation

Del-1 is an anti-inflammatory factor under various conditions. To determine whether Del-1 also plays an anti-inflammatory effect in prion pathogenesis, we analyzed microglial activation and astrogliosis in prion-infected terminally sick Del-1^+/+^, Del-1^+/−^ and Del-1^−/−^ mice. Histology failed to reveal overt differences in Iba1 (microglial marker) and glial fibrillary acidic protein (GFAP, astrocytic marker) immunoreactivity between Del-1^+/+^, Del-1^+/−^ and Del-1^−/−^ mice (Figure 5A-B). We then performed Western blots to detect Iba1 and GFAP protein in prion-infected terminally sick Del-1^+/+^, Del-1^+/−^ and Del-1^−/−^ mouse brains. Again, we observed similar levels of Iba1 and GFAP in all three groups (Figure 5C-D). These results suggest that, in contrast to other models of neurological disease such as EAE, Del-1 deficiency does not overtly affect prion-induced microglial activation and astrogliosis.

**Figure 5:**
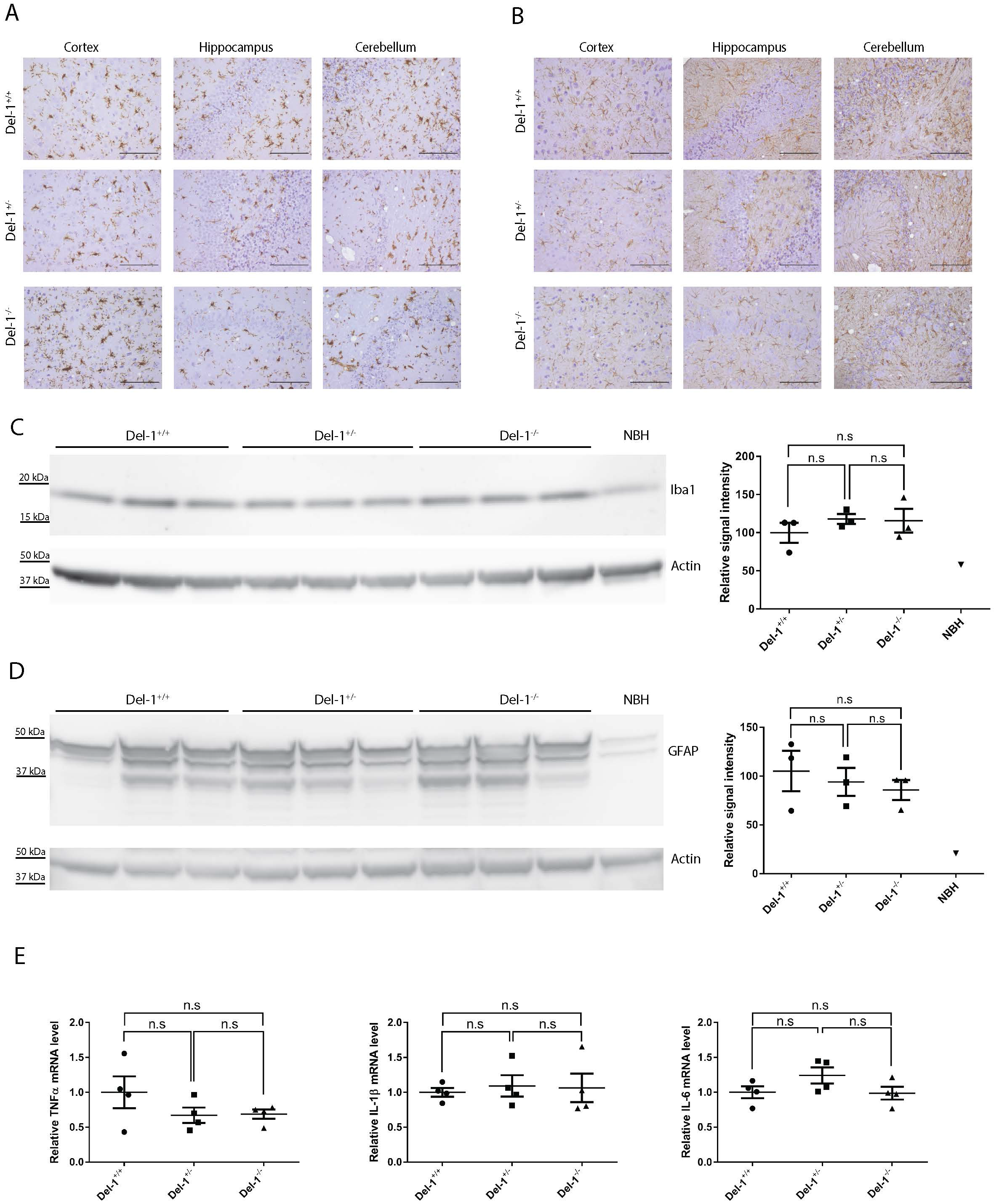
(A-B) Representative immunohistochemical staining for Iba1 (A) and GFAP (B) on cortex, hippocampus and cerebellum of terminally sick mouse brains from Del-1^+/+^ (WT), Del-1^+/−^ and Del-1^−/−^ littermates, Scale bars: 100um. (C) Left: Western blot for Iba1 in terminally sick mouse brains. Right: densitometric quantification of the Western blot revealed no significant difference of Iba1 levels between Del-1^+/+^ (WT), Del-1^+/−^ and Del-1^−/−^ littermates (n=3, n.s *p*>0.05). (D) Left: Western blot for GFAP in terminally sick mouse brains. Right: densitometric quantification of the Western blot showed no significant difference of GFAP levels between Del-1^+/+^ (WT), Del-1^+/−^ and Del-1^−/−^ littermates (n=3, n.s *p*>0.05). (E) qRT-PCR of cytokines TNFα, IL-6 and IL-1β expression revealed similar expression levels of these cytokines in terminally sick Del-1^+/+^ (WT), Del-1^+/−^ and Del-1^−/−^ littermates (n=4, n.s *p*>0.05).

We also performed cytokine profiling using qRT-PCR to assess the expression of proinflammatory cytokines including TNFα, IL-1β and IL6 in prion-infected terminally sick Del-1^+/+^, Del-1^+/−^ and Del-1^−/−^ mouse brains. This experiment did not detect obvious differences in cytokine expression between three groups (Figure 5E), suggesting that Del-1 deficiency fails to alter prion-induced cytokine expression.

## 4. Discussion

Neuroinflammation characterized by astrogliosis and microglial activation is considered a hallmark of various neurodegenerative conditions including Alzheimer’s disease and prion diseases (Aguzzi, et al., 2013a). A flurry of microglia-related genes has turned up in genome-wide association screens as risk factors of neurodegenerative disorders, suggesting that microglia may play a central role in these diseases (Baker, et al., 2006,Cruts, et al., 2006,Guerreiro, et al., 2013,Hollingworth, et al., 2011,Jonsson, et al., 2013,Naj, et al., 2011). Elucidating the molecular mechanisms by which microglia contributes to the pathogenesis of neurodegenerative conditions represents an important research frontier in contemporary neuroscience.

As the principal innate immune cells in the CNS, microglia serve as important defenders upon challenges. In mouse models of prion disease, microglia are found to play an overall neuroprotective role by clearing prions (Zhu, et al., 2016). Depletion or deficiency of microglia result in impaired prion clearance, enhanced PrP^Sc^ deposition and deteriorated prion pathogenesis. However, the molecular mechanisms underlying microglia-mediated prion clearance remain elusive (Aguzzi and Zhu, 2017). Mfge8, an opsonin that facilitates apoptotic cell phagocytosis, has been found to contribute to prion clearance. However, this effect is mouse strain-dependent (Kranich, et al., 2010), indicating that other Mfge8-associated molecules may complement this clearing function under certain conditions. Therefore, in this study we aimed to investigate the role of Del-1, a structural and functional homologue of Mfge8, in prion pathogenesis.

Del-1, initially detected in embryonic endothelial cells, is expressed in a developmentally restricted manner (Hidai, et al., 1998). Del-1 is also expressed by fetal thymic and liver macrophages (Hanayama, et al., 2004), as well as in the adrenal gland (Kanczkowski, et al., 2013). However, the expression pattern of Del-1 in adult mice remained to be delineated. We therefore characterized the expression of Del-1 in mice, focusing on the brain, because Del-1 expression level is highest in the brain, about three times greater than in the lungs, and only barely detectable in other organs. Interestingly, the cerebral expression of Del-1 is age-dependent showing an increase during the first four months followed by a stabilization. Del-1 expression also displays an age-dependent manner in other organs. In the gingiva, Del-1 expression decreases during aging (Eskan, et al., 2012).

To determine which cell types in the brain express Del-1, we co-stained β-galactosidase with various neural cell markers. We found that the primary cell types that express Del-1 in the brain are neurons. Although Del-1 shares structural and functional similarities to Mfge8, their expression patterns differ significantly. In fetal mice, thioglycolate-elicited macrophages express Mfge8 but not Del-1, whereas thymic and liver macrophages express Del-1 but not Mfge8 (Hanayama, et al., 2004). Intriguingly, this observation also pertains to the brain, Del-1 is expressed by neurons, whereas Mfge8 is known to be expressed by astrocytes (Kranich, et al., 2010).

The expression pattern of Del-1 in adults suggests that Del-1 not only plays a role during early development but also functions in adult in brain, lungs and adrenal glands. In the adrenal glands and lungs, Del-1 prevents leukocytes from adhering to endothelium and causing inflammation. In fact, Del-1^−/−^ mice demonstrated augmented inflammation within lungs and adrenal glands after a systemic inflammatory induction through the administration of lipopolysaccharide (Choi, et al., 2008,Kanczkowski, et al., 2013). The reduction of Del-1 expression in gingiva is associated with periodontitis (Eskan, et al., 2012). Those findings led to an assumption that Del-1 may represent a negative regulator for inflammation in general (Kanczkowski, et al., 2013). Therefore, we speculate that Del-1 may also act as anti-inflammatory factor in the brain. Indeed, Del-1 shows anti-inflammatory function in a mouse model of EAE (Choi, et al., 2015).

After defining the expression pattern and function of Del-1, we aimed to study whether the neuron-expressed Del-1 plays phagocytosis-promoting and/or anti-inflammation functions in prion diseases. Especially we sought to determine whether Del-1 complements Mfge8 in prion clearance in mice with a C57BL/6J genetic background, in which Mfge8 deficiency does not influence prion pathogenesis. After prion infection, we observed that Del-1^−/−^ mice experienced a similar incubation time compared with Del-1^+/−^ and wild type littermates, suggesting that Del-1 deficiency does not affect prion disease progression. More importantly, the PrP^Sc^ level and the prion seeding activity were not detectably affected by Del-1 deficiency, indicating that Del-1 is not a major contributor to prion clearance. Moreover, prion-induced neuroinflammation including astrocytosis and microglial activation was not altered in Del-1^−/−^ mouse brains, suggesting that Del-1 deficiency does not overtly affect prion-induced neuroinflammation.

In conclusion, our study revealed that Del-1 is expressed mainly by neurons in the mouse brain, and Del-1 expression is age-dependent. In the prion inoculation experiment, we demonstrated that Del-1 deficiency neither affects prion disease progression, nor lesion pattern, nor alters prion clearance or PrP^Sc^ deposition. Additionally, Del-1 deficiency does not affect prion-induced neuroinflammation. Therefore, neuron-expressed Del-1 is not a major determinant of prion pathogenesis and does not complement Mfge8 in prion clearance in mice with the C57BL/6J genetic background. Further studies are required to identify the molecular mechanisms underlying microglial uptake and clearance of prions.

## Acknowledgements

We thank the team of the Institute of Neuropathology, University Hospital Zurich, and in particular M. Delic, K. Arroyo, R. Moos, I. Abakumova and M. König for technical assistance. We thank Elisabeth J. Rushing for reading and editing the manuscript.We also gratefully thank Triantafyllos Chavakis from University Clinic Carl-Gustav-Carus Dresden University of Technology for providing us with Del-1^−/−^ mice. A. Aguzzi is the recipient of an Advanced Grant of the European Research Council (ERC, No. 250356) and is supported by grants from the European Union (PRIORITY, NEURINOX), the Swiss National Foundation (SNF, including a Sinergia grant), the Swiss Initiative in Systems Biology, SystemsX.ch (PrionX, SynucleiX), and the Klinische Forschungsschwerpunkte (KFSPs) “small RNAs” and “Human Hemato-Lymphatic Diseases”. The funders had no role in study design, data collection and analysis, decision to publish, or preparation of the manuscript. The authors declare no competing financial interests.

---

**Supplementary figure 1**: Western blots using anti-Del-1 antibodies that could not detect specific Del-1 band on brains collected from wild type mice. Del-1^−/−^ mouse brains were used as negative control.

**Supplementary figure 2**: Immunofluorescent co-staining of β-galactosidase and cell markers (GFAP, CD68 and MAG) on Del-1^−/−^ mouse brains. No colocalization was observed between β-galactosidase stain (green dots) and GFAP, CD68 or MAG, Scale bars: 10um.

## References

Aguzzi, A., Barres, B.A., Bennett, M.L. 2013a. Microglia: scapegoat, saboteur, or something else? Science 339(6116), 156–61. doi:10.1126/science.1227901.

Aguzzi, A., Nuvolone, M., Zhu, C. 2013b. The immunobiology of prion diseases. Nature reviews Immunology 13(12), 888–902. doi:10.1038/nri3553.

Aguzzi, A., Zhu, C. 2012. Five questions on prion diseases. PLoS pathogens 8(5), e1002651. doi:10.1371/journal.ppat.1002651.

Aguzzi, A., Zhu, C. 2017. Microglia in prion diseases. J Clin Invest. doi:10.1172/JCI90605.

Baker, M., Mackenzie, I.R., Pickering-Brown, S.M., Gass, J., Rademakers, R., Lindholm, C., Snowden, J., Adamson, J., Sadovnick, A.D., Rollinson, S., Cannon, A., Dwosh, E., Neary, D., Melquist, S., Richardson, A., Dickson, D., Berger, Z., Eriksen, J., Robinson, T., Zehr, C., Dickey, C.A., Crook, R., McGowan, E., Mann, D., Boeve, B., Feldman, H., Hutton, M. 2006. Mutations in progranulin cause tau-negative frontotemporal dementia linked to chromosome 17. Nature 442(7105), 916–9. doi:10.1038/nature05016.

Choi, E.Y., Chavakis, E., Czabanka, M.A., Langer, H.F., Fraemohs, L., Economopoulou, M., Kundu, R.K., Orlandi, A., Zheng, Y.Y., Prieto, D.A., Ballantyne, C.M., Constant, S.L., Aird, W.C., Papayannopoulou, T., Gahmberg, C.G., Udey, M.C., Vajkoczy, P., Quertermous, T., Dimmeler, S., Weber, C., Chavakis, T. 2008. Del-1, an endogenous leukocyte-endothelial adhesion inhibitor, limits inflammatory cell recruitment. Science 322(5904), 1101–4. doi:10.1126/science.1165218.

Choi, E.Y., Lim, J.H., Neuwirth, A., Economopoulou, M., Chatzigeorgiou, A., Chung, K.J., Bittner, S., Lee, S.H., Langer, H., Samus, M., Kim, H., Cho, G.S., Ziemssen, T., Bdeir, K., Chavakis, E., Koh, J.Y., Boon, L., Hosur, K., Bornstein, S.R., Meuth, S.G., Hajishengallis, G., Chavakis, T. 2015. Developmental endothelial locus-1 is a homeostatic factor in the central nervous system limiting neuroinflammation and demyelination. Mol Psychiatry 20(7), 880–8. doi:10.1038/mp.2014.146.

Cruts, M., Gijselinck, I., van der Zee, J., Engelborghs, S., Wils, H., Pirici, D., Rademakers, R., Vandenberghe, R., Dermaut, B., Martin, J.J., van Duijn, C., Peeters, K., Sciot, R., Santens, P., De Pooter, T., Mattheijssens, M., Van den Broeck, M., Cuijt, I., Vennekens, K., De Deyn, P.P., Kumar-Singh, S., Van Broeckhoven, C. 2006. Null mutations in progranulin cause ubiquitin-positive frontotemporal dementia linked to chromosome 17q21. Nature 442(7105), 920–4. doi:10.1038/nature05017.

Eskan, M.A., Jotwani, R., Abe, T., Chmelar, J., Lim, J.H., Liang, S., Ciero, P.A., Krauss, J.L., Li, F., Rauner, M., Hofbauer, L.C., Choi, E.Y., Chung, K.J., Hashim, A., Curtis, M.A., Chavakis, T., Hajishengallis, G. 2012. The leukocyte integrin antagonist Del-1 inhibits IL-17-mediated inflammatory bone loss. Nat Immunol 13(5), 465–73. doi:10.1038/ni.2260.

Falsig, J., Julius, C., Margalith, I., Schwarz, P., Heppner, F.L., Aguzzi, A. 2008. A versatile prion replication assay in organotypic brain slices. Nature neuroscience 11(1), 109–17. doi:10.1038/nn2028.

Frontzek, K., Pfammatter, M., Sorce, S., Senatore, A., Schwarz, P., Moos, R., Frauenknecht, K., Hornemann, S., Aguzzi, A. 2016. Neurotoxic Antibodies against the Prion Protein Do Not Trigger Prion Replication. PLoS One 11(9), e0163601. doi:10.1371/journal.pone.0163601.

Guerreiro, R., Wojtas, A., Bras, J., Carrasquillo, M., Rogaeva, E., Majounie, E., Cruchaga, C., Sassi, C., Kauwe, J.S., Younkin, S., Hazrati, L., Collinge, J., Pocock, J., Lashley, T., Williams, J., Lambert, J.C., Amouyel, P., Goate, A., Rademakers, R., Morgan, K., Powell, J., St George-Hyslop, P., Singleton, A., Hardy, J., Alzheimer Genetic Analysis, G. 2013. TREM2 variants in Alzheimer’s disease. The New England journal of medicine 368(2), 117–27. doi:10.1056/NEJMoa1211851.

Hanayama, R., Tanaka, M., Miwa, K., Nagata, S. 2004. Expression of developmental endothelial locus-1 in a subset of macrophages for engulfment of apoptotic cells. Journal of immunology 172(6), 3876–82.

Hidai, C., Zupancic, T., Penta, K., Mikhail, A., Kawana, M., Quertermous, E.E., Aoka, Y., Fukagawa, M., Matsui, Y., Platika, D., Auerbach, R., Hogan, B.L., Snodgrass, R., Quertermous, T. 1998. Cloning and characterization of developmental endothelial locus-1: an embryonic endothelial cell protein that binds the alphavbeta3 integrin receptor. Genes Dev 12(1), 21–33.

Hollingworth, P., Harold, D., Sims, R., Gerrish, A., Lambert, J.C., Carrasquillo, M.M., Abraham, R., Hamshere, M.L., Pahwa, J.S., Moskvina, V., Dowzell, K., Jones, N., Stretton, A., Thomas, C., Richards, A., Ivanov, D., Widdowson, C., Chapman, J., Lovestone, S., Powell, J., Proitsi, P., Lupton, M.K., Brayne, C., Rubinsztein, D.C., Gill, M., Lawlor, B., Lynch, A., Brown, K.S., Passmore, P.A., Craig, D., McGuinness, B., Todd, S., Holmes, C., Mann, D., Smith, A.D., Beaumont, H., Warden, D., Wilcock, G., Love, S., Kehoe, P.G., Hooper, N.M., Vardy, E.R., Hardy, J., Mead, S., Fox, N.C., Rossor, M., Collinge, J., Maier, W., Jessen, F., Ruther, E., Schurmann, B., Heun, R., Kolsch, H., van den Bussche, H., Heuser, I., Kornhuber, J., Wiltfang, J., Dichgans, M., Frolich, L., Hampel, H., Gallacher, J., Hull, M., Rujescu, D., Giegling, I., Goate, A.M., Kauwe, J.S., Cruchaga, C., Nowotny, P., Morris, J.C., Mayo, K., Sleegers, K., Bettens, K., Engelborghs, S., De Deyn, P.P., Van Broeckhoven, C., Livingston, G., Bass, N.J., Gurling, H., McQuillin, A., Gwilliam, R., Deloukas, P., Al-Chalabi, A., Shaw, C.E., Tsolaki, M., Singleton, A.B., Guerreiro, R., Muhleisen, T.W., Nothen, M.M., Moebus, S., Jockel, K.H., Klopp, N., Wichmann, H.E., Pankratz, V.S., Sando, S.B., Aasly, J.O., Barcikowska, M., Wszolek, Z.K., Dickson, D.W., Graff-Radford, N.R., Petersen, R.C., Alzheimer’s Disease Neuroimaging, I., van Duijn, C.M., Breteler, M.M., Ikram, M.A., DeStefano, A.L., Fitzpatrick, A.L., Lopez, O., Launer, L.J., Seshadri, S., consortium, C., Berr, C., Campion, D., Epelbaum, J., Dartigues, J.F., Tzourio, C., Alperovitch, A., Lathrop, M., consortium, E., Feulner, T.M., Friedrich, P., Riehle, C., Krawczak, M., Schreiber, S., Mayhaus, M., Nicolhaus, S., Wagenpfeil, S., Steinberg, S., Stefansson, H., Stefansson, K., Snaedal, J., Bjornsson, S., Jonsson, P.V., Chouraki, V., Genier-Boley, B., Hiltunen, M., Soininen, H., Combarros, O., Zelenika, D., Delepine, M., Bullido, M.J., Pasquier, F., Mateo, I., Frank-Garcia, A., Porcellini, E., Hanon, O., Coto, E., Alvarez, V., Bosco, P., Siciliano, G., Mancuso, M., Panza, F., Solfrizzi, V., Nacmias, B., Sorbi, S., Bossu, P., Piccardi, P., Arosio, B., Annoni, G., Seripa, D., Pilotto, A., Scarpini, E., Galimberti, D., Brice, A., Hannequin, D., Licastro, F., Jones, L., Holmans, P.A., Jonsson, T., Riemenschneider, M., Morgan, K., Younkin, S.G., Owen, M.J., O’Donovan, M., Amouyel, P., Williams, J. 2011. Common variants at ABCA7, MS4A6A/MS4A4E, EPHA1, CD33 and CD2AP are associated with Alzheimer’s disease. Nat Genet 43(5), 429–35. doi:10.1038/ng.803.

Jonsson, T., Stefansson, H., Steinberg, S., Jonsdottir, I., Jonsson, P.V., Snaedal, J., Bjornsson, S., Huttenlocher, J., Levey, A.I., Lah, J.J., Rujescu, D., Hampel, H., Giegling, I., Andreassen, O.A., Engedal, K., Ulstein, I., Djurovic, S., Ibrahim-Verbaas, C., Hofman, A., Ikram, M.A., van Duijn, C.M., Thorsteinsdottir, U., Kong, A., Stefansson, K. 2013. Variant of TREM2 associated with the risk of Alzheimer’s disease. The New England journal of medicine 368(2), 107–16. doi:10.1056/NEJMoa1211103.

Kanczkowski, W., Chatzigeorgiou, A., Grossklaus, S., Sprott, D., Bornstein, S.R., Chavakis, T. 2013. Role of the endothelial-derived endogenous anti-inflammatory factor Del-1 in inflammation-mediated adrenal gland dysfunction. Endocrinology 154(3), 1181–9. doi:10.1210/en.2012-1617.

Klotzsche-von Ameln, A., Cremer, S., Hoffmann, J., Schuster, P., Khedr, S., Korovina, I., Troullinaki, M., Neuwirth, A., Sprott, D., Chatzigeorgiou, A., Economopoulou, M., Orlandi, A., Hain, A., Zeiher, A.M., Deussen, A., Hajishengallis, G., Dimmeler, S., Chavakis, T., Chavakis, E. 2017. Endogenous developmental endothelial locus-1 limits ischaemia-related angiogenesis by blocking inflammation. Thromb Haemost 117(6), 1150–63. doi:10.1160/TH16-05-0354.

Kranich, J., Krautler, N.J., Falsig, J., Ballmer, B., Li, S., Hutter, G., Schwarz, P., Moos, R., Julius, C., Miele, G., Aguzzi, A. 2010. Engulfment of cerebral apoptotic bodies controls the course of prion disease in a mouse strain-dependent manner. The Journal of experimental medicine 207(10), 2271–81. doi:10.1084/jem.20092401.

Kranich, J., Krautler, N.J., Heinen, E., Polymenidou, M., Bridel, C., Schildknecht, A., Huber, C., Kosco-Vilbois, M.H., Zinkernagel, R., Miele, G., Aguzzi, A. 2008. Follicular dendritic cells control engulfment of apoptotic bodies by secreting Mfge8. The Journal of experimental medicine 205(6), 1293–302. doi:10.1084/jem.20071019.

Maekawa, T., Hosur, K., Abe, T., Kantarci, A., Ziogas, A., Wang, B., Van Dyke, T.E., Chavakis, T., Hajishengallis, G. 2015. Antagonistic effects of IL-17 and D-resolvins on endothelial Del-1 expression through a GSK-3beta-C/EBPbeta pathway. Nat Commun 6, 8272. doi:10.1038/ncomms9272.

Mitroulis, I., Chen, L.S., Singh, R.P., Kourtzelis, I., Economopoulou, M., Kajikawa, T., Troullinaki, M., Ziogas, A., Ruppova, K., Hosur, K., Maekawa, T., Wang, B., Subramanian, P., Tonn, T., Verginis, P., von Bonin, M., Wobus, M., Bornhauser, M., Grinenko, T., Di Scala, M., Hidalgo, A., Wielockx, B., Hajishengallis, G., Chavakis, T. 2017. Secreted protein Del-1 regulates myelopoiesis in the hematopoietic stem cell niche. J Clin Invest 127(10), 3624–39. doi:10.1172/JCI92571.

Mitroulis, I., Kang, Y.Y., Gahmberg, C.G., Siegert, G., Hajishengallis, G., Chavakis, T., Choi, E.Y. 2014. Developmental endothelial locus-1 attenuates complement-dependent phagocytosis through inhibition of Mac-1-integrin. Thromb Haemost 111(5), 1004–6. doi:10.1160/TH13-09-0794.

Naj, A.C., Jun, G., Beecham, G.W., Wang, L.S., Vardarajan, B.N., Buros, J., Gallins, P.J., Buxbaum, J.D., Jarvik, G.P., Crane, P.K., Larson, E.B., Bird, T.D., Boeve, B.F., Graff-Radford, N.R., De Jager, P.L., Evans, D., Schneider, J.A., Carrasquillo, M.M., Ertekin-Taner, N., Younkin, S.G., Cruchaga, C., Kauwe, J.S., Nowotny, P., Kramer, P., Hardy, J., Huentelman, M.J., Myers, A.J., Barmada, M.M., Demirci, F.Y., Baldwin, C.T., Green, R.C., Rogaeva, E., St George-Hyslop, P., Arnold, S.E., Barber, R., Beach, T., Bigio, E.H., Bowen, J.D., Boxer, A., Burke, J.R., Cairns, N.J., Carlson, C.S., Carney, R.M., Carroll, S.L., Chui, H.C., Clark, D.G., Corneveaux, J., Cotman, C.W., Cummings, J.L., DeCarli, C., DeKosky, S.T., Diaz-Arrastia, R., Dick, M., Dickson, D.W., Ellis, W.G., Faber, K.M., Fallon, K.B., Farlow, M.R., Ferris, S., Frosch, M.P., Galasko, D.R., Ganguli, M., Gearing, M., Geschwind, D.H., Ghetti, B., Gilbert, J.R., Gilman, S., Giordani, B., Glass, J.D., Growdon, J.H., Hamilton, R.L., Harrell, L.E., Head, E., Honig, L.S., Hulette, C.M., Hyman, B.T., Jicha, G.A., Jin, L.W., Johnson, N., Karlawish, J., Karydas, A., Kaye, J.A., Kim, R., Koo, E.H., Kowall, N.W., Lah, J.J., Levey, A.I., Lieberman, A.P., Lopez, O.L., Mack, W.J., Marson, D.C., Martiniuk, F., Mash, D.C., Masliah, E., McCormick, W.C., McCurry, S.M., McDavid, A.N., McKee, A.C., Mesulam, M., Miller, B.L., Miller, C.A., Miller, J.W., Parisi, J.E., Perl, D.P., Peskind, E., Petersen, R.C., Poon, W.W., Quinn, J.F., Rajbhandary, R.A., Raskind, M., Reisberg, B., Ringman, J.M., Roberson, E.D., Rosenberg, R.N., Sano, M., Schneider, L.S., Seeley, W., Shelanski, M.L., Slifer, M.A., Smith, C.D., Sonnen, J.A., Spina, S., Stern, R.A., Tanzi, R.E., Trojanowski, J.Q., Troncoso, J.C., Van Deerlin, V.M., Vinters, H.V., Vonsattel, J.P., Weintraub, S., Welsh-Bohmer, K.A., Williamson, J., Woltjer, R.L., Cantwell, L.B., Dombroski, B.A., Beekly, D., Lunetta, K.L., Martin, E.R., Kamboh, M.I., Saykin, A.J., Reiman, E.M., Bennett, D.A., Morris, J.C., Montine, T.J., Goate, A.M., Blacker, D., Tsuang, D.W., Hakonarson, H., Kukull, W.A., Foroud, T.M., Haines, J.L., Mayeux, R., Pericak-Vance, M.A., Farrer, L.A., Schellenberg, G.D. 2011. Common variants at MS4A4/MS4A6E, CD2AP, CD33 and EPHA1 are associated with late-onset Alzheimer’s disease. Nat Genet 43(5), 436–41. doi:10.1038/ng.801.

Sailer, A., Bueler, H., Fischer, M., Aguzzi, A., Weissmann, C. 1994. No propagation of prions in mice devoid of PrP. Cell 77(7), 967–8.

Shin, J., Maekawa, T., Abe, T., Hajishengallis, E., Hosur, K., Pyaram, K., Mitroulis, I., Chavakis, T., Hajishengallis, G. 2015. DEL-1 restrains osteoclastogenesis and inhibits inflammatory bone loss in nonhuman primates. Science translational medicine 7(307), 307ra155. doi:10.1126/scitranslmed.aac5380.

Takahashi, K., Rochford, C.D., Neumann, H. 2005. Clearance of apoptotic neurons without inflammation by microglial triggering receptor expressed on myeloid cells-2. The Journal of experimental medicine 201(4), 647–57. doi:10.1084/jem.20041611.

Wang, Y., Cella, M., Mallinson, K., Ulrich, J.D., Young, K.L., Robinette, M.L., Gilfillan, S., Krishnan, G.M., Sudhakar, S., Zinselmeyer, B.H., Holtzman, D.M., Cirrito, J.R., Colonna, M. 2015. TREM2 lipid sensing sustains the microglial response in an Alzheimer’s disease model. Cell 160(6), 1061–71. doi:10.1016/j.cell.2015.01.049.

Zhu, C., Herrmann, U.S., Falsig, J., Abakumova, I., Nuvolone, M., Schwarz, P., Frauenknecht, K., Rushing, E.J., Aguzzi, A. 2016. A neuroprotective role for microglia in prion diseases. The Journal of experimental medicine 213(6), 1047–59. doi:10.1084/jem.20151000.

Zhu, C., Herrmann, U.S., Li, B., Abakumova, I., Moos, R., Schwarz, P., Rushing, E.J., Colonna, M., Aguzzi, A. 2015a. Triggering receptor expressed on myeloid cells-2 is involved in prion-induced microglial activation but does not contribute to prion pathogenesis in mouse brains. Neurobiology of aging 36(5), 1994–2003. doi:10.1016/j.neurobiolaging.2015.02.019.

Zhu, C., Schwarz, P., Abakumova, I., Aguzzi, A. 2015b. Unaltered Prion Pathogenesis in a Mouse Model of High-Fat Diet-Induced Insulin Resistance. PLoS One 10(12), e0144983. doi:10.1371/journal.pone.0144983.

